# Structure Model Analysis Of Phosphorylation Dependent Binding And Sequestration Of SARS-COV-2 Encoded Nucleocapsid Protein By Protein 14-3-3

**DOI:** 10.1101/2020.09.16.299362

**Authors:** Pierre Limtung, H.Y. Lim Tung

**Author notes:** Correspondence to: H.Y. Lim Tung, Peptide and Protein Chemistry Research Laboratory, Nacbraht Biomedical Research Institute, 3164 21st Street, Suite 122, Astoria (NYC), NY 11106, USA, E mail.

## Abstract

Phosphorylation of serines 197 and 206 of SARS-COV-2 Nucleocapsid protein (NCp) enhanced the stability and binding efficiency and sequestration of NCp to Protein 14-3-3 by increasing the Stability Energy (ΔGstability energy) and Binding Energy (ΔΔGbinding energy) from ~545 Kcal/mol to ~616 Kcal/mol, and from 108 Kcal/mol to ~228 Kcal/mol respectively. The calculated Binding Energy Difference (ΔΔGbinding energy difference) between dephospho-NCp-14-3-3 complex and phospho-NCp-13-3-3 complex was ~72 Kcal/mol. Phosphorylations of serines 186, 197, 202 and 206, and threonines 198 and 205 NCp also caused an increase in the Stability Energy (ΔGstability energy) and Binding Energy (ΔΔGbinding energy) from ~545 Kcal/mol to ~617, 616, 583, 580, 574, 564 and 566 Kcal/mol and from ~108 Kcal/mol to ~228, 216, 184, 188, 184, 174 and 112 Kcal/mol respectively. Phosphorylation of NCp on serines 197 and 206 caused a decrease in Stability Energy and Binding Energy from ~698 Kcal/mol to 688 Kcal/mol, and from ~91 Kcal/mol to ~82 Kcal/mol for the dimerization of NCp. These results support the existence of a phosphorylation dependent cellular mechanism to bind and sequester NCp.

## Introduction

SARS-COV-2, the etiologic agent of COVID-19 [1–4] enters its host cells via binding to a cellular receptor identified as ACE2 [5–8]. The viability, infectivity and virulence of SARS-COV-2 are dependent upon successful replication, transcription and packaging of SARS-COV-2 genome. The molecular mechanisms that underlie controls of replication, transcription and packaging of SARS-COV-2 genome are not well understood [9–11]. An essential controller of the replication, transcription and packaging of SARS-COV-2 genome is the Nucleocapsid protein (NCp). It has been proposed that NCp plays an essential role as a co-factor in the initiation and control of the replication, transcription and packaging of the SARS-COV-2 genome [10–20]. Dimerization of NCp is a prerequisite step in the replication, transcription and packaging of the SARS-COV-2 genome [12–20]. Inhibiting the functions of NCp represents an attractive way to control SARS-COV-2 viability, infections and virulence [11, 21–24]. There is also evidence that NCp may be an attractive antigen molecule for vaccine development purpose [25–29]. It has been suggested that there is a cellular response mechanism for preventing dimerization of NCp which involves the phosphorylation dependent binding and sequestration of NCp by Protein 14-3-3 [30]. A phosphorylation rich domain of NCp containing at least two binding sites (RNpSTP and RGTpSP) for Protein 14-3-3 located in the linker region of NCp which is exposed at the surface of NCp has been identified [30]. Through structure model analysis, it was demonstrated that only monomeric NCp phosphorylated on serines 197 and and 206 can fit in the binding groove of Protein 14-3-3 [30].

Phosphorylation of serines 197 and 206 within the motifs RNpSTP and RGTpSP of NCp by C-TAK-1 has been proposed to constitute the formation of the binding and sequestration sites for Protein 14-3-3 [30]. Other phosphorylations sites, including serines 186, 198, 202 and 205 are also present in the phosphorylation rich domain of NCp. Serines 186, 197 and 202 of NCp become mutated to phenylalanine, leucine and asparagine in SARS-COV-2 strains/sub-strains isolated from individuals from Iran, Spain and India respectively [30]. The significance of these mutations is not known. It has been argued that mutations of phospho-sites 186, 197 and 206 allows NCp to avoid its sequestration by Protein 14-3-3 [30]. The importance of phosphorylation of serines 186, 197, 198, 202, 205 and 206 remain to be established. Phosphorylation of NCp by various Protein kinases, including GSK-3, Cdk1 and PKA have been described and shown to affect the location and function of NCp in the cell [30–34]. Here, we use computerized structure model analysis and thermodynamic calculation to determine the roles of multi-sites phosphorylation in the binding and sequestration of NCp by Protein 14-3-3. The results presented here support the conclusion that cells infected with SARS-COV-2 possess a phosphorylation dependent cellular response mechanism for the binding and sequestration of NCp by Protein 14-3-3 involving phosphorylations of serines 186, 197, 198, 202, 205 and 206.

## Methods

The structure of dephospho-SARS-COV-2 Nucleocapsid protein (NCp) was rendered de novo using the Quark Program pursuant to et Zu and Zhang [35, 36]. Phosphorylation of NCp was performed using the Build Model Program of FoldX pursuant to Guerois et al. and Schymkowitz et al. [37, 38]. Docking experiments to identify the binding and sequestration of NCp by Protein 14-3-3 and dimerization of NCp were performed using the ZDOCK pursuant to Pierce et al. [39]. NCps rendered in this work and Protein 14-3-3 (1YZ5) based on the structure determination of Benzinger et al. [40] were analyzed and visualized by the CCP4 Molecular Graphics Program Version 2.10.11 as described by Mc Nicolas et al [41] and the ZMM Molecular Modeling Program as described by Garden and Zhorov [42].

Determination and calculation of Stability Energy (ΔG_stability energy_) of the protein complexes and non-complexed proteins was performed using the Stability Program of FoldX as described by Guerois et al. and Schymkowitz et al. [37, 38]. Binding Energy (ΔΔG_binding energy_) of protein complex was determined and calculated using the Analyze Complex Program of FoldX as described by Guerois et al. and Schymkowitz et al. [37, 38]. Binding Energy Difference (ΔΔΔG_binding enrgy difference_) between dephospho-NCp-Protein 14-3-3 complex and phospho-NCp-Protein 14-3-3 complex was calculated pursuant to Teng et al. and Nishi et al. [43–45] from the equation: ΔΔΔG_binding energy difference_ = ΔΔG_binding energy_ (binding energy of phospho-NCp-14-3-3 complex) - ΔΔG_binding energy_ (binding energy of dephospho-NCp-14-3-3 complex.

The above formula was also used to calculate the Binding Energy Difference (ΔΔΔG_binding energy_) between dephospho-NCp Dimer and phospho-NCp Dimer.

## Results

The results of de novo rendering of the structure of dephospho NCp (amino acids 123-310) which encompasses the phosphorylation rich domain of NCp (amino acids 185-209) using the Quark Program pursuant to Xu and Zhang [35, 36] was previously described [30]. Figure 1 is a summary of the docking experiments that identified the interaction between unphosphorylated NCp and Protein 14-3-3. Unphosphorylated NCp did not form a stable complex with Protein 14-3-3 and did not fit itself in the binding groove of Protein 14-3-3.

**Figure 1.**
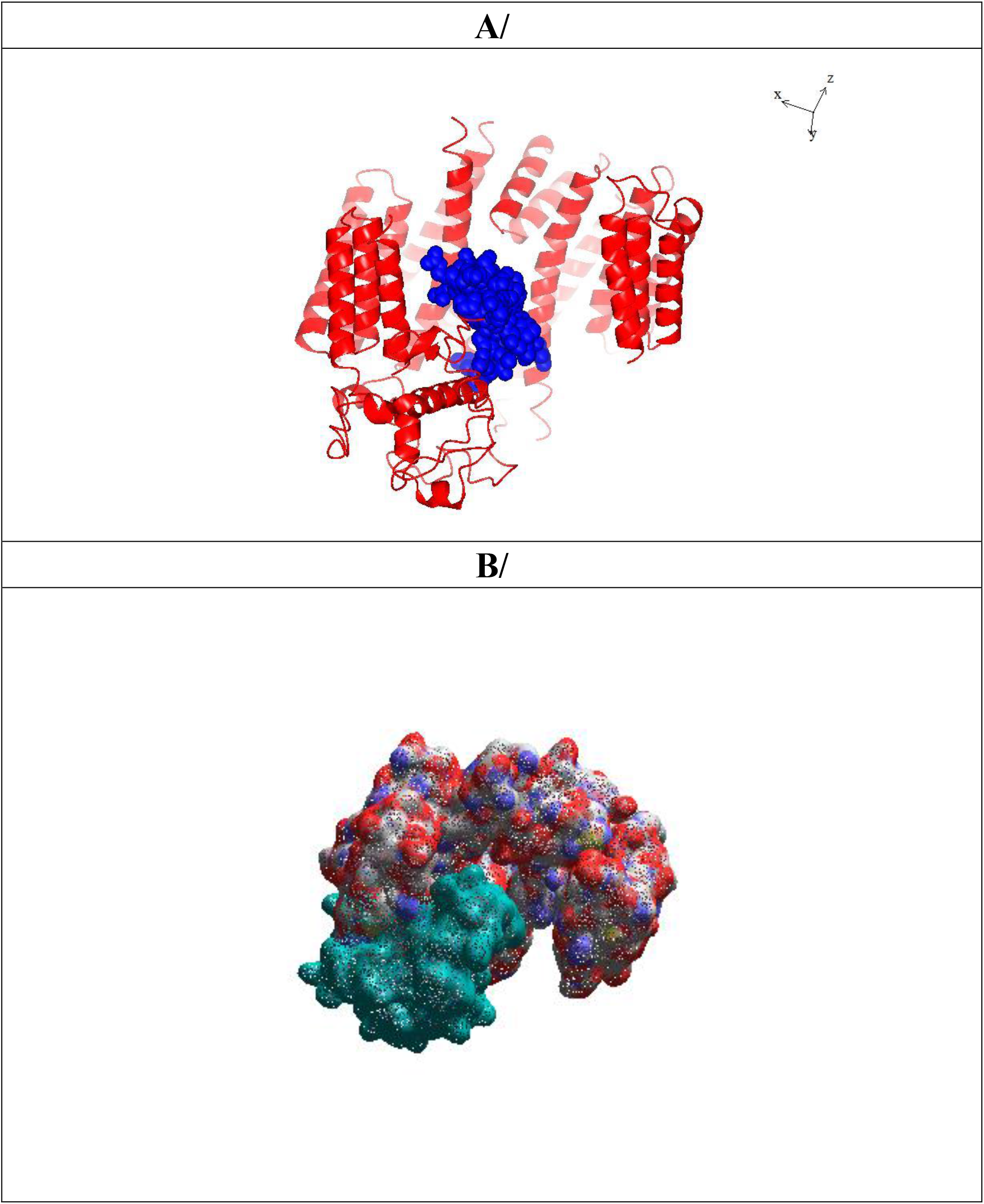
A: Ribbon structure (red) of dephosphorylated SARS-COV-2 Nucleocapsid protein (NCp) and Protein 14-3-3 pre-complex, rendered as described in Method Section. The amino acid sequence of the phosphorylation rich domain (blue spheres) is. B: Realistic rendering of dephosphorylated SARS-COV-2 Nucleocapsid protein (NCp) (Cyan) and Protein 14-3-3 pre-complex (Red and Blue).

Figure 2 summarizes the docking experiments which showed that NCp phosphorylated on serine 197 and serine 206 fitted neatly into the binding groove of Protein 14-3-3 and also formed a stable complex with Protein 14-3-3. The results of Computerized Thermodynamic Calculations are summarized in Table 1. The Stability Energy (ΔG_stability_) for the dephosphorylated NCp-Protein 14-3-3 and NCp phosphorylated on serines 197 and 206-Protein 14-3-3 complexes were calculated to be ~545 Kcal/mol and ~616 Kcal/mol respectively. The Binding Energy (ΔΔG_binding_) for the interactions between the components of dephosphorylated NCp-Protein 14-3-3 and NCp phosphorylated on serines 197 and 206-Protein 14-3-3 complexes were determined to be ~108 Kcal/mol and ~228 Kcal/mol respectively. These results indicated that phosphorylation of NCp on serines 197 and 206 resulted in enhanced stability and binding affinity of the NCp-Protein 14-3-3 complex. That phosphorylation on NCp on serine 197 and 206 enhanced the stability and binding affinity of NCp-Protein 14-3-3 complex was confirmed from the calculated Binding Energy Difference (ΔΔΔG_binding energy difference_) (~120 Kcal/mol) (Table 1). Pursuant to Teng et al. [42–44] a positive value for Binding Energy Difference (ΔΔΔG_binding energy difference_) indicates an enhancement of stability and binding affinity of the protein components in the protein complex as compared to the unmodified protein complex. Phosphorylation of serine 197 was accompanied by an increase in Binding Energy (G) from ~108 Kcal/mol to 184 Kcal/mol while phosphorylation of serine 206 was associated with a small increase of Binding Energy from ~108 Kcal/mol to ~112 Kcal/mol. On the other hand, upon phosphorylations of both serines 197 and 206, the Binding Energy (G) was ~228 Kcal/mol. The Binding Energy Differences of ~76 Kcal/mol following phosphorylation of serine 197, ~4 Kcal/mol following phosphorylation of 206 and ~120 following phosphorylations of both serines 197 and 206 suggest that the effects of the phosphorylations of serine 197 and 206 within the motifs which are recognized and phosphorylated by C-TAK1 were not additive but most likely synergistic.

**Figure 2.**
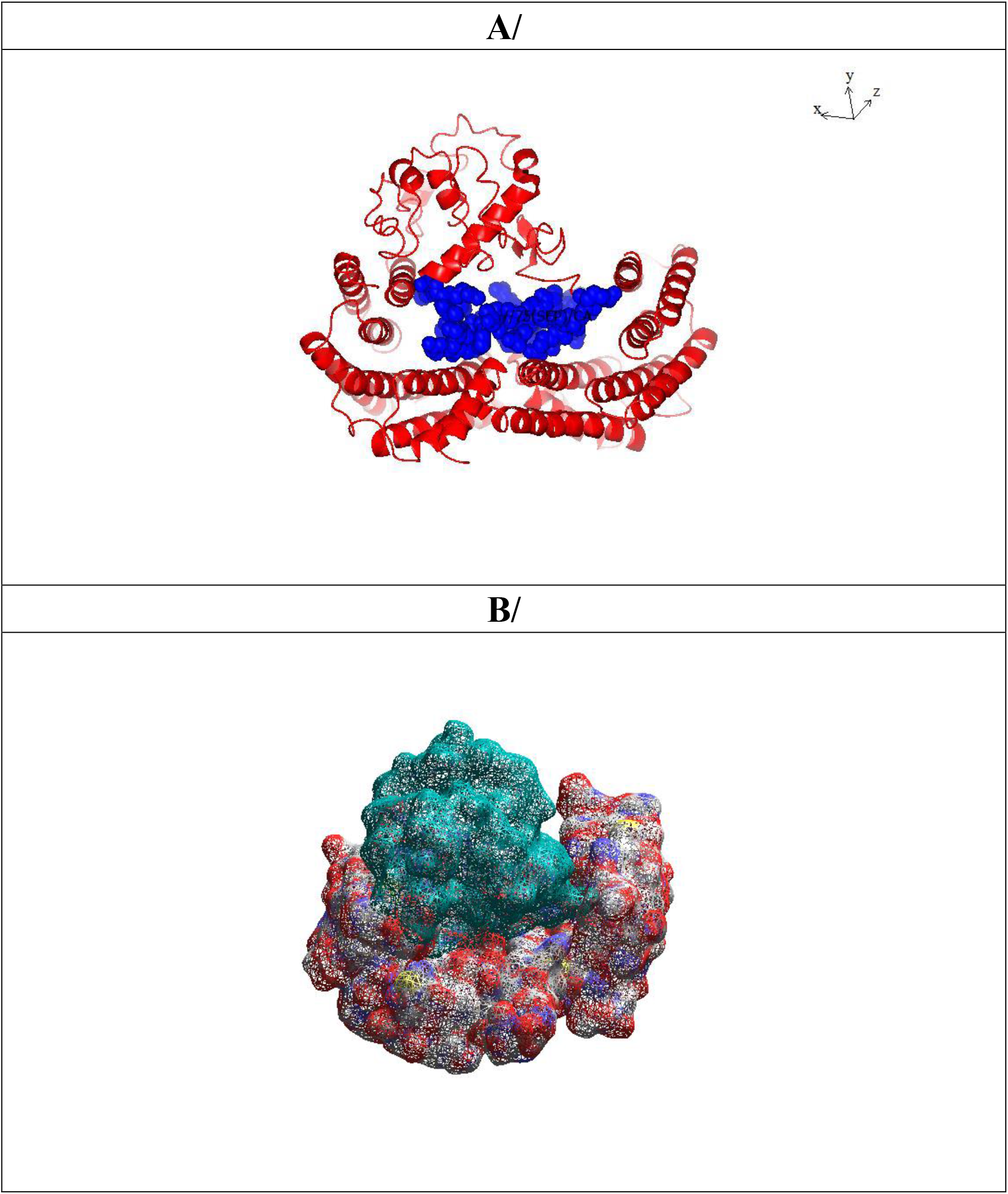
A: Ribbon structure (red) of SARS-COV-2 Nucleocapsid protein (NCp) phosphorylated on serines 197 and 206 and Protein 14-3-3 complex, rendered as described in Method Section. Blue spheres represent the phosphorylation rich domain, B: Realistic rendering SARS-COV-2 Nucleocapsid protein (NCp (cyan) phosphorylated on serines 197 and 206 and Protein 14-3-3(red and blue) complex.

**Table 1:** Thermodynamic calculations, including Stability (ΔG_stability_), Binding Energy (ΔΔG_binding energy)_ and Binding Energy Difference (ΔΔΔG_binding energy difference_) that underlie the sequestration of dephospho- and various phospho-NCp-Protein 14-3-3 complexes.

| | Stability<br>Energy<br>($\Delta G$)<br>Kcal/mol | Binding<br>Energy<br>($\Delta\Delta G$)<br>Kcal/mol | Binding Energy<br>Difference<br>($\Delta\Delta\Delta G$)<br>Kcal/mol |
| --- | --- | --- | --- |
| Dephospho-NCp-14-3-3 complex | ~545 | ~108 | ~000 |
| Phospho-serines 197/206-NCp-14-3-3 complex | ~617 | ~228 | ~120 |
| Phospho-serine 186-NCp-14-3-3 complex | ~616 | ~216 | ~107 |
| Phospho-serine 197-NCp-14-3-3 complex | ~583 | ~184 | ~76 |
| Phospho-threonine 198-NCp-14-3-3 complex | ~580 | ~188 | ~80 |
| Phospho-serine 202-NCp-14-3-3 complex | ~574 | ~184 | ~76 |
| Phospho-threonine 205-NCp-14-3-3 complex | ~564 | ~174 | ~66 |
| Phospho-serine 206-NCp-14-3-3 complex | ~566 | ~112 | ~4 |

Phosphorylations of serines 186, 197, 202 and 206, and threonines 198 and 205 of NCp were all accompanied by an increase of the Stability Energy (ΔG_stability energy_) from ~545 Kcal/mol to ~616, 583, 580, 574 and 566 Kcal/mol. Phosphorylations of serines 186, 197, 202, and 206, and threonines 198 and 205 of NCp were also all associated with increase of Binding Energy (ΔΔG_binding energy_) from ~108 Kcal/mol to ~216, 184, 188, 184, 174 and 112 Kcal/mol respectively. The Binding Energy Difference (ΔΔΔG_binding energy difference_) between dephosphorylated NCp and NCp phosphorylated on serines 186, 197, 202 and 206, and threonines 198 and 205 were calculated to be ~107, 76, 80, 76, 66 and 4 Kcal/mol respectively. These results indicate that phosphorylation of serines 186, 197, 202 and 206, and threonines 198 and 205 of NCp were all accompanied by enhanced stability and binding affinity of the protein components of the NCp-Protein 14-3-3 complexes.

Figure 3 summarizes the docking experiments which showed the dimerization of unphosphorylated NCp. The Stability Energy (ΔG_stability energy_) of the unphosphorylated NCp dimer and Binding Energy (ΔΔG_binding energy_) of the interaction between NCp molecules of the NCp dimer were calculated to be ~698 Kcal/mol and ~91 Kcal/mol respectively. Figure 4 depicts the docking experiments for the dimerization of NCp phosphorylated on serines 197 and 206. The Stability Energy ((ΔG_stability energy_) of the unphosphorylated NCp dimer and Binding Energy (ΔΔG_binding energy_) of the interaction between NCp molecules of the NCp dimer were calculated to be ~688 Kcal/mol and ~82 Kcal/mol respectively. The Binding Energy Difference (ΔΔΔG_binding energy difference_) between dephospho-NCp dimer and NCp phosphorylated on serines 197 and 206 was calculated to be ~ −10 Kcal/mol (Table 2). Pursuant to Teng et al. and Nishi et al. [43–45], negative Binding Energy Difference (ΔΔΔG_binding energy difference_) indicates reductions of stability and binding affinity of the protein components in the protein complex as compared to the unmodified protein complex. Except for phosphorylation of serine 197 which did not produce a significant change in the Stability Energy of the Ncp-Protein 14-3-3 complex, the phosphorylations of serines 186, 202 and 206, and threonines 198 and 205 were all accompanied by a decrease in Binding Stability (ΔG) from ~698 Kcal/mol to ~596, 693, 675, 678 and 664 Kcal/mol. There were significant decreases of Binding Energy (ΔΔG) from ~91 Kcal/mol to ~63, 87, 87, 74, 79 and 61 Kcal/mol when serines 186, 197, 202 and 206, and threonines 198 and 205 were phosphorylated (Table 2). There were also significant decreases in Binding Energy Difference (Table 2). These results are consistent with the conclusion that phosphorylations of NCp caused a decrease in the stability and binding affinity of the two components of the NCp dimer.

**Figure 3.**
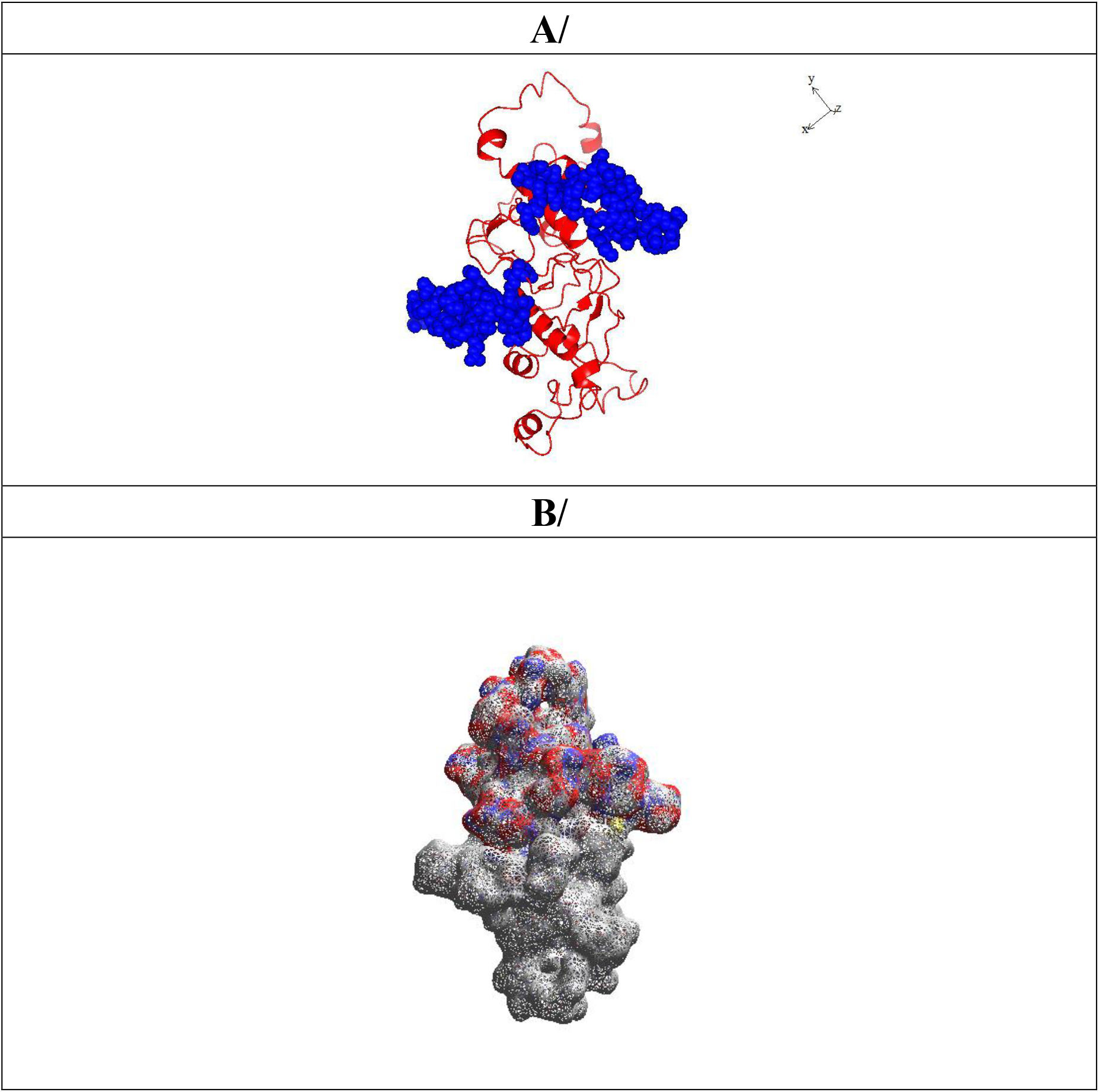
A: Ribbon structure (red) of unphosphorylated SARS-COV-2 Nucleocapsid protein (NCp) dimer. Blue spheres represent the phosphorylation rich domain of NCp), rendered as described in Method Section. B: Realistic rendering of dimer of unphosphorylated SARS-COV-2 Nucleocapsid protein (NCp).

**Figure 4.**
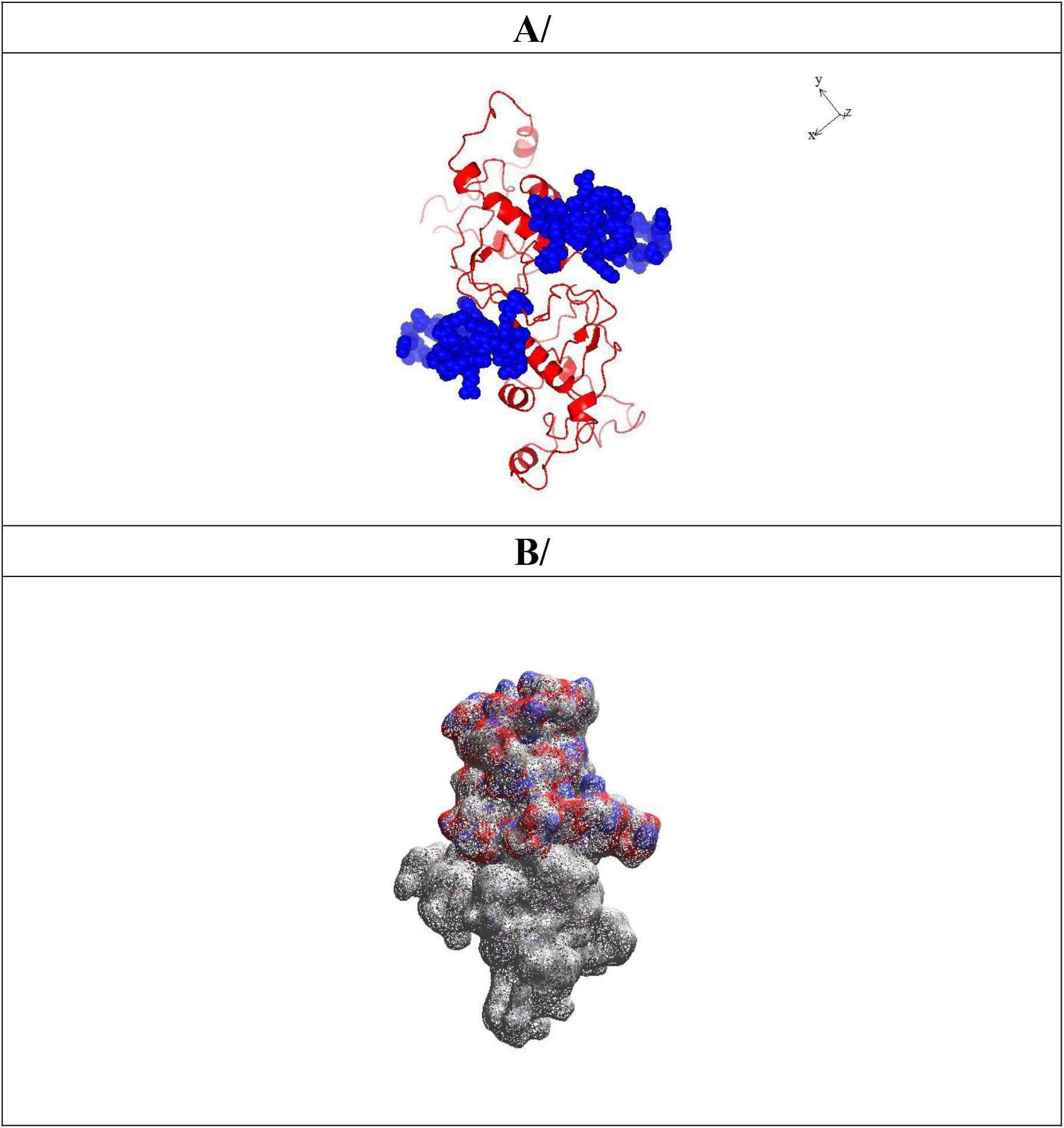
A: Ribbon structure (red) of phosphorylated SARS-COV-2 Nucleocapsid protein (NCp) dimer. Blue spheres represent the phosphorylation rich domain of NCp), rendered as described in Method Section. B: Realistic rendering of dimer of unphosphorylated SARS-COV-2 Nucleocapsid protein (NCp).

**Table 2:** Thermodynamic calculations, including Stability (ΔG_stability_), Binding Energy (ΔΔG_binding energy)_ and Binding Energy Difference (ΔΔΔG_binding energy difference_) that underlie the dimerization of dephospho- and various phospho-NCps.

| | Stability<br>Energy<br>($\Delta G$)<br>Kcal/mol | Binding<br>Energy<br>($\Delta\Delta G$)<br>Kcal/mol | Binding Energy<br>Difference<br>($\Delta\Delta\Delta G$)<br>Kcal/mol |
| --- | --- | --- | --- |
| Dephospho-NCp-NCp complex | ~698 | ~91 | ~ 00 |
| Phospho-serines 197/206-NCp-NCp complex | ~688 | ~82 | ~ -10 |
| Phospho-serine 186-NCp-NCp complex | ~596 | ~63 | ~ -28 |
| Phospho-serine 197-NCp-NCp complex | ~699 | ~87 | ~ -4 |
| Phospho-threonine 198-NCp-NCp complex | ~693 | ~87 | ~ -4 |
| Phospho-serine 202-NCp-NCp complex | ~675 | ~74 | ~ -17 |
| Phospho-threonine 205-NCp-NCp complex | ~678 | ~79 | ~ -12 |
| Phospho-serine 206-NCp-NCp complex | ~664 | ~61 | ~ -30 |

## Discussion

NCp is an essential co-factor in the replication, transcription and packaging of the SARS-COV-2 genome [9–20]. Inhibiting the function of NCp is therefore an attractive approach to prevent the transmission, infection and virulence of SARS-COV-2 [21–29]. Dimerization and oligomerization of NCp are essential for its function as a co-factor of replication, transcription and packaging of the SARS-COV-2 genome [13, 15–17, 19]. We have previously shown that Protein 14-3-3 can bind and sequester monomeric NCp in a phosphorylation dependent manner [30]. We showed that within NCp, there is a phosphorylation rich domain located between the dimerization and RNA binding domains that harbors two phosphorylation sites, serines 197 and 206 that are phosphorylated by C-TAK1 [30]. The amino acid sequence around phospho-serines 197 and 206 form binding and sequestration sites (RNpSTP and RGTpSP) for Protein 14-3-3 which is an important signaling molecule involved in the control of the cell cycle, cell survival and cell death [30].

In addition to serines 197 and 206, there are other phosphorylation sites in the phosphorylation rich domain of NCp, including serine 186 which can be phosphorylated CKI, serine 198 which can be phosphorylated by the proline dependent protein kinases, including GSK-3 and Cdk1, serine 202 which can be phosphorylated by GSK-3 and serine 205 which can be phosphorylated by PKA. While it is fairly obvious that phospho-serines 197 and 206 together with their surrounding amino acids act as binding and sequestration sites for Protein 14-3-3 [30, 46–49], the roles of phospho-serine 186, 198, 202 and 206 were not. In the present study, we have determined that phosphorylating serines 197 and 206 of NCp caused the latter to enter the binding groove of Protein 14-3-3 and to be bound and sequestered by Protein 14-3-3. Through computerized Structure Model Analysis and Thermodynamic Calculation, we have demonstrated that phosphorylation of serines 197 and 206 caused an increase in the Stability Energy (ΔG_stability energy_) and Binding Energy (ΔΔG_binding energy_) of NCp phosphorylated on serine 197 and 206-Protein 14-3-3 complex. We have also shown that phosphorylations of serines 186, 198, 202 and 205 of NCp were also accompanied by an increase in Stability Energy (ΔG_stability energy_) and Binding Energy (ΔΔG_binding energy_) of NCp-Protein 14-3-3 complex. The calculated Binding Energy Difference (ΔΔG_binding energy difference_) confirmed that there was an enhancement of stability and binding affinity of NCp-Protein 14-3-3 complex associated with phosphorylation of NCp on serines 186, 198, 202 and 205 of NCp.

On the other hand, we showed that phosphorylation of serines 197 and 206 caused a decrease in the Stability Energy (ΔG_stability energy_) and Binding Energy (ΔΔG_binding energy_) of the dimerization of NCp dimer. Calculation of the Binding Energy Difference (ΔΔG_binding energy difference_) between dephospho NCp dimer and phospho-NCp dimer confirmed such assessment. He et al. [16] have reported that the phosphorylation rich domain of NCp was important in its oligomerization. Peng et al. [17] have reported that phosphorylation of NCp within the phosphorylation rich domain impaired its oligomerization. Klann et al. [50] have described the phosphorylations of several sites in NCp. However, their functional significance remains to be clarified. Here, we showed that phosphorylation of serines 186, 197 and 206 of NCp was responsible for the inhibition of NCp dimerization. In support of the conclusions of He et al. [16] and Peng et al. [17], the present work which used Computerized Structure Model Analysis and Thermodynamic Calculation provides evidence that the phosphorylation rich domain of NCp was involved in preventing the dimerization of NCp.

The results of the present work suggest two complementary mechanisms of phosphorylation dependent inhibition of NCp functions: (i) Phosphorylation dependent binding and sequestration of NCp by Protein 14-3-3 involving phosphorylation of serines 197 and 206 within the motifs RNpSTP and RGTpSP which are phosphorylated by C-TAK-1 and recognized by Protein 14-3-3 [46–49], and also of serines 186, 198, 202 and 205 which are phopshorylated by CKI, GSK-3/Cdk1, GSK-3 and PKA respectively [29], and (ii) Phosphorylation dependent inhibition of NCp dimerization involving phosphorylation of serine 186 and serines 197 and 206 [present work]. In view of the fact that dimerization of NCp is necessary for it to act as an essential co-factor for the replication, transcription and packaging of SARS-COV-2 genome [9–20], it is proposed that sequestration of NCp by Protein 14-3-3 is a cellular response mechanism for inhibiting replication, transcription and packaging of the SARS-COV-2 genome exists in the host cells of SARS-COV-2. Inhibitors of NCp dimerization would include activators of the protein kinases which phosphorylate serines 186, 197, 198, 202, 205 and 206 of NCp or molecules that directly prevent NCp dimerization by acting at the interface between the NCp monomers or at remote positions.

## Acknowledgements

This work was supported by the Nacbraht Biomedical Research Institute Fund.

## Author contributions

H.Y. Lim Tung came up with the concept and the questions, performed the experiments with Pierre Limtung, analyzed the results with Pierre Limtung and wrote the paper with Pierre Limtung.

Pierre Limtung performed the experiments with H.Y. Lim Tung, analyzed the results with H.Y. Lim Tung and wrote the paper with Pierre Limtung.

## References

1. Zhu, N., Zhang, D., Wang, W., Li, X., Yang, B., Song, J., Zhao, X., Huang, B., Shi, W., Lu, R., Niu, P., Zhan, F., Ma, X., Wang, D., Xu, W., Wu, G., Gao, G.F. and Tan, W. (2019) N. Engl. J. Med., Vol. 382, pp 727–733. A novel coronavirus from patients with pneumonia in China, 2019.

2. Lu, R., Zhao, X., Li, J., Niu, P., Yang, B., Wu, H., Wang, W., Song, H., Huang, B., Zhu, N., Bi, Y., Ma, X., Zhan, F., Wang, L., Hu, T., Zhou, H., Hu, Z., Zhou, W., Zhao, L., Chen, J., Meng, Y., Wang, J., Lin, Y., Yuan, J., Xie, Z., Ma, J., Liu, W.J., Wang, D., Xu, W., Holmes, E.C., Gao, G.F., Wu, G., Chen, W., Shi, W. and Tan, W. (2020) Lancet, Vol. 395, pp 565–574. Genomic characterisation and epidemiology of 2019 novel coronavirus: implications for virus origins and receptor binding.

3. Zhou, P., Yang, X.L., Wang, X.G., Hu, B., Zhang, L., Zhang, W., Si, H.R., Zhu, Y., Li, B., Huang, C.L., Chen, H.D., Chen, J., Luo, Y., Guo, H., Jiang, R.D., Liu, M.Q., Zhao, K., Chen, Q.J., Deng, F., Liu, L.L., Yan, B., Zhan, Q.J., Deng, F., Liu, L.L., Yan, B., Zhan, F.X., Wang, Y.Y., Gao, G.F. and Shi, Z.L. (2020) Nature, Vol. 579, pp 270–273. A pneumonia outbreak associated with a new coronavirus of probable bat origin.

4. Wu, F., Zhao, S., Yu, B., Chen, Y.-M., Wang, W., Song, Z.-G, Hu, Y., Tao, Z.-W., Tian, J.-H., Pei, Y.-Y., Yuan, M.-L., Zhang, Y.-L., Dai, F.-H., Liu, Y., Wang, Q.-M., Zheng, J.-J., Xu, L., Holmes, E.C. and Zhang, Y.Z. (2020) Nature, Vol., 579, pp 265–269. A new coronavirus associated with human respiratory disease in China.

5. Li, W., Moore, M.J. N, Vasilieva, N., Sui, J. Wong, S.K., Berne MA, Somasundaran, M., Sullivan, J.L., Luzuriaga, K., Greenough, T.C., Choe, H. and Farzan, M. (2003) Nature, Vol. 426, pp 450–453. Angiotensin-converting enzyme 2 is a functional receptor for the SARS coronavirus.

6. Li, F., Li, W., Farzan, M. and Harrison, S.C. (2005) Science, Vol. 309, pp 1864–1868. Structure of SARS coronavirus spike receptor-binding domain complexed with receptor.

7. Kuba, K., Imai, Y., Rao, S., Guo, F., Guan, B., Huan, Y., Yang, P., Zhang, Y., Deng, W., Bao, L., Zhang, B., Liu, G., Wang, Chappell, Liu, Y., Zheng, D., Leibbrandt, A., Wada, T., Slutsky, A., Liu, D., Qin, C., Jiang, C. and Penninger, J.M. (2005) Nat. Med., Vol. 11, pp 875–879. A crucial role of angiotensin converting enzyme 2 (ACE2) in SARS coronavirus-induced lung injury.

8. Shang, J., Ye, G., Shi, K., Wan, Y., Luo, C., Aihara, H., Geng, Q., Auerbach and Li, F. (2020) Nature, Vol. 581, pp 221–224. Structural basis of receptor recognition by SARS-CoV-2.

9. Stertz, S., Reichelt, M., Spiegel, M., Kuri, T., Martínez-Sobrido, L., García-Sastre, A., Weber, F. and Kochs, G. (2007). Virology, Vol. 361, pp 304–315. The intracellular sites of early replication and budding of SARS coronavirus.

10. Verheije, M. H., Hagemeijer, M. C., Ulasli, M., Reggiori, F., Rottier, P. J., Masters, P. S., and de Haan, C. A. (2010) J. Virol., Vol. 84, pp 11575–11579. The coronavirus nucleocapsid protein is dynamically associated with the replication-transcription complexes.

11. Cong, Y., Ulasli, M., Schepers, H., Mauthe, M., V’kovski, P., Kriegenburg, F., Thiel, V., de Haan, C.A.M. and Reggiori, F. (2020) J. Virol., DOI: 10.1128/JVI.01925-19. Nucleocapsid Protein Recruitment to Replication-Transcription Complexes Plays a Crucial Role in Coronaviral Life Cycle.

12. Tylor, S., Andonove, A., Cutts, T., Cao, J., Grudesky, E., Van Domselaar, G., Li, X. and He, R. (2009) Can. J. Microbiol., Vol. 55, pp 254–260. The SR-rich motif in SARS-CoV nucleocapsid protein is important for virus replication.

13. Chen, C. Y., Chang, C. K., Chang, Y. W., Sue, S. C., Bai, H. I., Riang, L., Hsiao, C. D., and Huang, T. H. (2007) J. Mol. Biol., 10.1016/j.jmb.2007.02.069. Structure of the SARS coronavirus nucleocapsid protein RNA-binding dimerization domain suggests a mechanism for helical packaging of viral RNA.

14. Hsieh, P. K., Chang, S. C., Huang, C. C., Lee, T. T., Hsiao, C. W., Kou, Y. H., Chen, I. Y., Chang, C. K., Huang, T. H., & Chang, M. F. (2005) J. Virol., Vol. 79, pp 13848–13855. Assembly of severe acute respiratory syndrome coronavirus RNA packaging signal into virus-like particles is nucleocapsid dependent.

15. Yu, I.-M., Gustafson, C.L.T., Diao, J., Burgner II, J.W., Li, Z. and Chen, J. (2005) J. Biol. Chem., Vol. 280, pp 23280–23286. Recombinant Severe Acute Respiratory Syndrome (SARS) Coronavirus Nucleocapsid Protein Forms a Dimer through Its C-terminal Domain

16. He, R., Dobie, F., Ballantine, M., Leeson, A., Li, Y., Bastien, N., Cutts, T., Andonov, A., Cao, J., Booth, T.F., Plummer, F.A., Tyler, S., Baker, L. and Li, X. (2004) Biochem. Biophys. Res. Commun., Vol. 316, pp 476–483. Analysis of multimerization of the SARS coronavirus nucleocapsid protein.

17. Peng, T.-Y., Lee, K.-R. and Tarn, W.-Y. (2008) FEBS J., Vol. 275, pp 4152–4163. Phosphorylation of the arginine/serine dipeptide-rich motif of the severe acute respiratory syndrome coronavirus nucleocapsid protein modulates its multimerization, translation inhibitory activity and cellular localization.

18. Surjit, M., Liu, B., Kumar, P., Chow, V. T., & Lal, S. K. (2004) Biochem. Biophys. Res. Commun., Vol. 317, pp 1020–1036. The nucleocapsid protein of the SARS coronavirus is capable of self-association through a C-terminal 209 amino acid interaction domain.

19. Luo, H., Ye, F., Sun, T., Yue, L., Peng, S., Chen, J., Li, G., Du, Y., Xie, Y., Yang, Y., Shen, J., Wang, Y., Shen, X., & Jiang, H. (2004) Biophys. Chem., Vol. 112, pp 15–25. In vitro biochemical and thermodynamic characterization of nucleocapsid protein of SARS.

20. Chang, C. K., Sue, S. C., Yu, T. H., Hsieh, C. M., Tsai, C. K., Chiang, Y. C., Lee, S. J., Hsiao, H. H., Wu, W. J., Chang, C. F., & Huang, T. H. (2005) FEBS Letters, Vol. 579, pp 5663–5668. The dimer interface of the SARS coronavirus nucleocapsid protein adapts a porcine respiratory and reproductive syndrome virus-like structure.

21. Gao, W., Tamin, A., Soloff, A., D’Aiuto, L., Nwanegbo, E., Robbins, P. D., Bellini, W. J., Barratt-Boyes, S., and Gambotto, A. (2003) Lancet, Vol. 362, pp 1895–1896. Effects of a SARS-associated coronavirus vaccine in monkeys.

22. Kang, S., Yang, M., Hong, Z., Zhang, L., Huang, Z., Chen, X., He, S., Zhou, Z., Zhou, Z., Chen, Q., Yan, Y., Zhang, X. S., Shan, H. and Chen, S. (2020) Acta Pharma. Sinica, Vol. 10, pp 1228–1238. Crystal structure of SARS-CoV-2 nucleocapsid protein RNA binding domain reveals potential unique drug targeting sites.

23. Mukherjee, D., and Upasana, R. (2020) chemRxiv, DOI: 10.26434/chemrxiv.12587336.v2. SARS-CoV-2 Nucleocapsid Assembly Inhibitors: Repurposing Antiviral and Antimicrobial Drugs Targeting Nucleocapsid-RNA Interaction.

24. Yadav, R., Imran, M., Dhamija, P., Kapil Suchal, K. and Handu, S. (2020) J. Biomol. Struc. Dynamics, DOI: 10.1080/07391102.2020.1778536. Virtual screening and dynamics of potential inhibitors targeting RNA binding domain of nucleocapsid phosphoprotein from SARS-CoV-2.

25. Liu, S. J., Leng, C. H., Lien, S. P., Chi, H. Y., Huang, C. Y., Lin, C. L., Lian, W. C., Chen, C. J., Hsieh, S. L., & Chong, P. (2006) Vaccine, Vol. 24, pp 3100–3108. Immunological characterizations of the nucleocapsid protein based SARS vaccine candidates.

26. Lin, Y., Shen, X., Yang, R. F., Li, Y. X., Ji, Y. Y., He, Y. Y., Shi, M. D., Lu, W., Shi, T. L., Wang, J., Wang, H. X., Jiang, H. L., Shen, J. H., Xie, Y. H., Wang, Y., Pei, G., Shen, B. F., Wu, J. R. and Sun, B. (2003). Cell Res., Vol. 13, pp 141–145. Identification of an epitope of SARS-coronavirus nucleocapsid protein.

27. Peng, H., Yang, L.-T., Wang, L.-Y., Li, J., Huang, J., Lu, Z-Q., Koup, R.A., Bailer, R.T. and Wu, C.-Y. (2006) Virology, Vol. 351, pp 466–475.

28. Kalita, P., Padhi, A. K., Zhang, K., & Tripathi, T. (2020) Microbial Pathogenesis, DOI: 10.1016/j.micpath.2020.104236 Design of a peptide-based subunit vaccine against novel coronavirus SARS-CoV-2.

29. Dutta, N.K., Mazumdar, K. and Gordy, J.T. (2020) J. Virol., DOI: 10.1128/JVI.00647-20. The Nucleocapsid Protein of SARS–CoV-2: a Target for Vaccine Development.

30. Tung, H.Y.L. and Limtung, Pierre (2020) Biochem. Biophys. Res. Commun., DOI: 10.1016/j.bbrc.2020.08.024. Mutations in the phosphorylation sites of SARS-CoV-2 encoded nucleocapsid protein and structure model of sequestration by protein 14-3-3.

31. Surjit, M., Kumar, R., Mishra, R. N., Reddy, M. K., Chow, V. T., and Lal, S. K. (2005) J. Virol., Vol. 79, pp 11476–11486. The severe acute respiratory syndrome coronavirus nucleocapsid protein is phosphorylated and localizes in the cytoplasm by 14-3-3-mediated translocation

32. C.H. Wu, S.-H. Yeh, Y.G. Tsay et al. Glycogen synthase kinase-3 regulates the phosphorylation of severe acute respiratory syndrome coronavirus nucleocapsid protein and viral replication. J. Biol. Chem. 284 (2009) 5229–52239.

33. C.H. Wu, P.J. Chen, S.-H Yeh, Nucleocapsid phosphorylation and RNA helicase DDX1 recruitment enables Coronavisrus transition from discontinuous transcription. Cell Host and Microbe 16 (2014) 462–472.

34. Carlson, C.R., Asfaha, J.B., Ghent, C.M., Howard, C.J., Hartooni, N. and Morgan, D.O. (2020) bioRxiv, DOI: 10.1101/2020.06.28.176248. Phosphorylation modulates liquid-liquid phase separation of the SARS-CoV-2 N protein.

35. D. Xu, Y. Zhang, Ab initio protein structure assembly using continuous structure fragments and optimized knowledge-based force field. (2012) Proteins 80 (2012) 1715–1735.

36. D. Xu, Y. Zhang, Toward optimal fragment generations for ab initio protein structure assembly. Proteins 81 (2013) 229–239.

37. Guerois, R., Nielson, J.E. and Serrano, L. (2002) J. Mol. Biol., Vol. 320, pp 369–387. Predicting changes in the stability of proteins and protein complexes: a study of more than 1000 mutations.

38. Joost Schymkowitz, J., Jesper Borg, J., Francois Stricher, F., Robby Nys, R., Frederic Rousseau, F. and Luis Serrano, L. (2005) Nucleic Acids Res., Vol. 33, pp W382–W388. The FoldX web server: an online force field.

39. Pierce, B.G., Wiehe, K., Hwang, H., Kim, B.H., Vreven, T. and Weng, Z. (2014) Bioinformatics, Vol. 30, pp 1771–1773. ZDOCK Server: Interactive Docking Prediction of Protein-Protein Complexes and Symmetric Multimers.

40. A. Benzinger, G.M. Popowicz, J.K. Joy et al. The crystal structure of the non-liganded 14-3-3sigma protein: insights into determinants of isoform specific ligand binding and dimerization. Cell Res. 15 (2005) 219–227.

41. McNicholas, S., Potterson, E., Wilson, K.S. and Noble, M.E.M. (2011) Acta Cryst., Vol. D67, pp 386–394. Presenting your structures: the CCP4mg molecular-graphics software.

42. Garden, D.P. and B.S. Zhorov, B.S. (2010) J. Comp. Aided Mol. Des., Vol. 25, pp 91–105. Docking flexible ligands in proteins with a solvent exposure-and distance-dependent dielectric function.

43. Teng, S., Srivastava, A. K., Schwartz, C. E., Alexov, E., & Wang, L. (2010) Int. J. Comput. Biol. Drug Des., Vol. 3, pp 334–349. Structural assessment of the effects of amino acid substitutions on protein stability and protein-protein interaction.

44. Nishi, H., Tyagi, M., Teng, S., Shoemaker, B.A., Hashimoto, K., Alexov, E., Wuchty, S. and Panchenko, A. R. (2013) Plos One, DOI: 10.1371/journal.pone.0066273. Cancer missense mutations alter binding properties of proteins and their interaction networks.

45. Nishi, H, Hashimoto, K. and Panchenko, A.R. (2011) Structure, Vol. 19, pp 1807–1815. Phosphorylation in protein-protein binding: effect on stability and function.

46. S. Ogg, B. Gabrielli, H. Piwnica-Worms, Purification of a Serine Kinase That Associates with and Phosphorylates Human Cdc25C on Serine 216. J. Biol. Chem. 269 (1994) 30461–30469.

47. J. Muller, D.A. Ritt, T.D. Copeland, D.K. Morrison (2003) EMBO J., Vol. 22, 4431–4442. Functional analysis of C-TAK1 substrate binding and identification of PKP2 as a new C-TAK1 substrate.

48. A.J. Muslin, J.W. Tanner, P.M. Allen et al. (1996) Cell, Vol. 84, pp 889–897. Interaction of 14-3-3 with signaling proteins is mediated by the recognition of phospho-serine.

49. M.B. Yaffe, K. Rittinger, S. Volinia, et al. (1997) Cell, Vol. 91, pp 961–971. The structural basis for 14-3-3:phosphopeptide binding specificity.

50. Klann, K., Bojkova, D., Tascher, G., Ciesek, S., Münch, C., & Cinatl, J. (2020) Mol. Cell. DOI: 10.1016/j.molcel.2020.08.006 Growth factor receptor signaling inhibition prevents SARS-CoV-2 replication.

